# Dimensional reduction of phenotypes from 53,000 mouse models reveals a diverse landscape of gene function

**DOI:** 10.1101/2021.06.10.447851

**Authors:** Tomasz Konopka, Letizia Vestito, Damian Smedley

## Abstract

Animal models have long been used to study gene function and the impact of genetic mutations on phenotype. Through the research efforts of thousands of research groups, systematic curation of published literature, and high-throughput phenotyping screens, the collective body of knowledge for the mouse now covers the majority of protein-coding genes. We here collected data for over 53,000 mouse models with mutations in over 15,000 genomic markers and characterized by more than 254,000 annotations using more than 9,000 distinct ontology terms. We investigated dimensional reduction and embedding techniques as means to facilitate access to this diverse and high-dimensional information. Our analyses provide the first visual maps of the landscape of mouse phenotypic diversity. We also summarize some of the difficulties in producing and interpreting embeddings of sparse phenotypic data. In particular, we show that data preprocessing, filtering, and encoding have as much impact on the final embeddings as the process of dimensional reduction. Nonetheless, techniques developed in the context of dimensional reduction create opportunities for explorative analysis of this large pool of public data, including for searching for mouse models suited to study human diseases.

## Introduction

Measuring the consequences of genetic mutations on organism-level phenotype is instrumental for describing gene function. It is a laborious process that requires breeding animals with controlled genotypes, performing a variety of assays, and describing phenotypes in a systematic fashion. For the mouse as a model organism, the collective knowledge of genotype-phenotype associations now covers around 15,000 genes (Blake et al., 2021). At the current pace of research, it may approach genome-wide coverage within a few years. A comprehensive phenomics dataset would impact many applications, for example, supporting efforts to identify the causes of rare genetic diseases (Justice & Dhillon, 2016; Meehan et al., 2017). However, there are currently few established methods to analyze phenomic data at scale, both for interactive exploration and for machine learning. Given recent advances in dimensional reduction, this promising approach may bring insight to mouse phenotype data and facilitate its integration with other omic datasets.

Phenotype data consist of links between the genetic characteristics of an animal model and sets of observations. For the mouse, the latter are usually tracked using the mammalian phenotype (MP) ontology (Smith & Eppig, 2015). The ontology is a collection of more than 13,000 concepts, also called MP terms, that are related through a hierarchy. For example, a phenotype describing ‘increased heart weight’ is a more precise annotation for the phenotype of ‘abnormal heart weight’, which in turn is a specific type of ‘cardiovascular system phenotype’. Individual terms in the ontology are thus not independent. However, despite the hierarchical connections, the space of possible phenotypic abnormalities is of high dimension. It covers all organ systems, fertility, and other factors, and animal models can be described by any combination of ontology terms.

High-dimensional data poses challenges both for explorative analysis and for machine learning (ML). The explorative analysis intends to place preliminary findings in context and to direct in-depth studies. It is often a manual process that relies on visualizations. Projection of large datasets into two dimensions is thus a common technique for this purpose; by placing data items in a scatter plot, it helps to convey similarity between many data items through their positions and relative distances in the embedding. Machine-learning models, by contrast, usually serve to automate decision making. In principle, they are not hindered by unintuitive data. However, training ML models requires fixing values for free parameters. When data is of high-dimension, ML models must estimate many more free parameters than with low-dimensional inputs. Using dimensional reduction may improve training efficiency, especially when there is limited data available.

A canonical approach to dimensional reduction of tabular data uses principal component analysis. It consists of a rotation - a linear transformation - followed by truncation to a fixed number of principal components, or dimensions, that incorporate the most variability. However, nonlinear embedding techniques can capture more relationships among the original data into the same number of dimensions. Examples include neural network auto-encoders (Hinton & Salakhutdinov, 2006), t-SNE (van der Maaten & Hinton, 2008), UMAP (Becht et al., 2019), EmbedSOM (Kratochvíl et al., 2019), PHATE (Moon et al. 2019), and Poincare maps (Klimovskaia et al. 2020). Approaches have also been implemented for non-tabular data. For example, node2vec provides embeddings of graphs (Grover & Leskovec, 2016). Importantly, recent implementations are computationally efficient and can process many thousands of data items. These methods have been instrumental in exploring atlases of mouse transcriptomic data (The Tabula Muris Consortium, 2018; Han, et al., 2018; Rosenberg et al., 2018; Saunders et al., 2018; Cao et al., 2019; Kalucka et al., 2020). They also open possibilities to analyze the phenotypic landscape of animal models.

Because ontology terms carry hierarchical relationships, dedicated methods have been proposed to embed the terms into low-dimensional spaces: Onto2vec (Smaili, et al., 2018), Opa2vec (Smaili et al., 2019), HiG2Vec (Kim et al., 2020), and Owl2vec (Chen et al., 2020). These approaches have also explored describing sets of ontology terms, particularly from the gene ontology. A common approach to embedding sets of ontology terms is through composition, or averaging, of the coordinates of the individual terms. This is motivated by observations in the context of word-based embeddings (Mikolov et al., 2013). It presents encouraging results also in the biological context, for example for the prediction of gene-gene interactions (Duong et al., 2020). However, linear composition conflicts with the nonlinearity inherent in dimensional reduction and is bound to lose effectiveness for sets with many terms. Moreover, ontology-based dimensional reduction techniques have not been explored in the context of animal phenotyping.

In this work, we examine a public dataset of mouse phenotypes and visualize the landscape of mouse phenotypic variation. We consider dimensional reduction approaches based on a range of mathematical techniques. We show that approaches that integrate sets of phenotypes into vectors or use text-based descriptions are more informative than approaches that rely on embeddings of individual ontology terms or on graphs. Yet, because mouse phenotype data is sparse, results can be sensitive to incomplete phenotyping and pre-processing steps.

## Results

### Embeddings of ontologies offer insight for small annotation sets

Ontology structures are relatively stable and evolve only slowly in time. Furthermore, the mammalian phenotype (MP) ontology that catalogs possible phenotypic aberrations is agnostic to animal species. It is thus natural to consider dimensional reduction of the space spanned by MP terms as a basis for visualizing phenotype data from the mouse and other model organisms. We produced embeddings of MP terms in two dimensions using two distinct approaches (Methods). In the first approach, we computed semantic similarities between text descriptions of MP terms and then created an embedding using uniform manifold approximation and projection (UMAP) (Becht et al., 2019). For comparison, we also constructed a graph of the hierarchical relations between MP terms and then created an embedding using node2vec (Grover & Leskovec, 2016). Of the two, the text-based approach placed the ontology root near the center, grouped ontology terms into small clusters, and produced a visual pattern that conveyed the multi-dimensionality of the ontology (Figure 1A). In contrast, the graph-based approach hid distinctions between phenotype domains (Figure S1). We continued to investigate animal models and diseases via the text-based embedding of MP terms.

**Figure 1.**
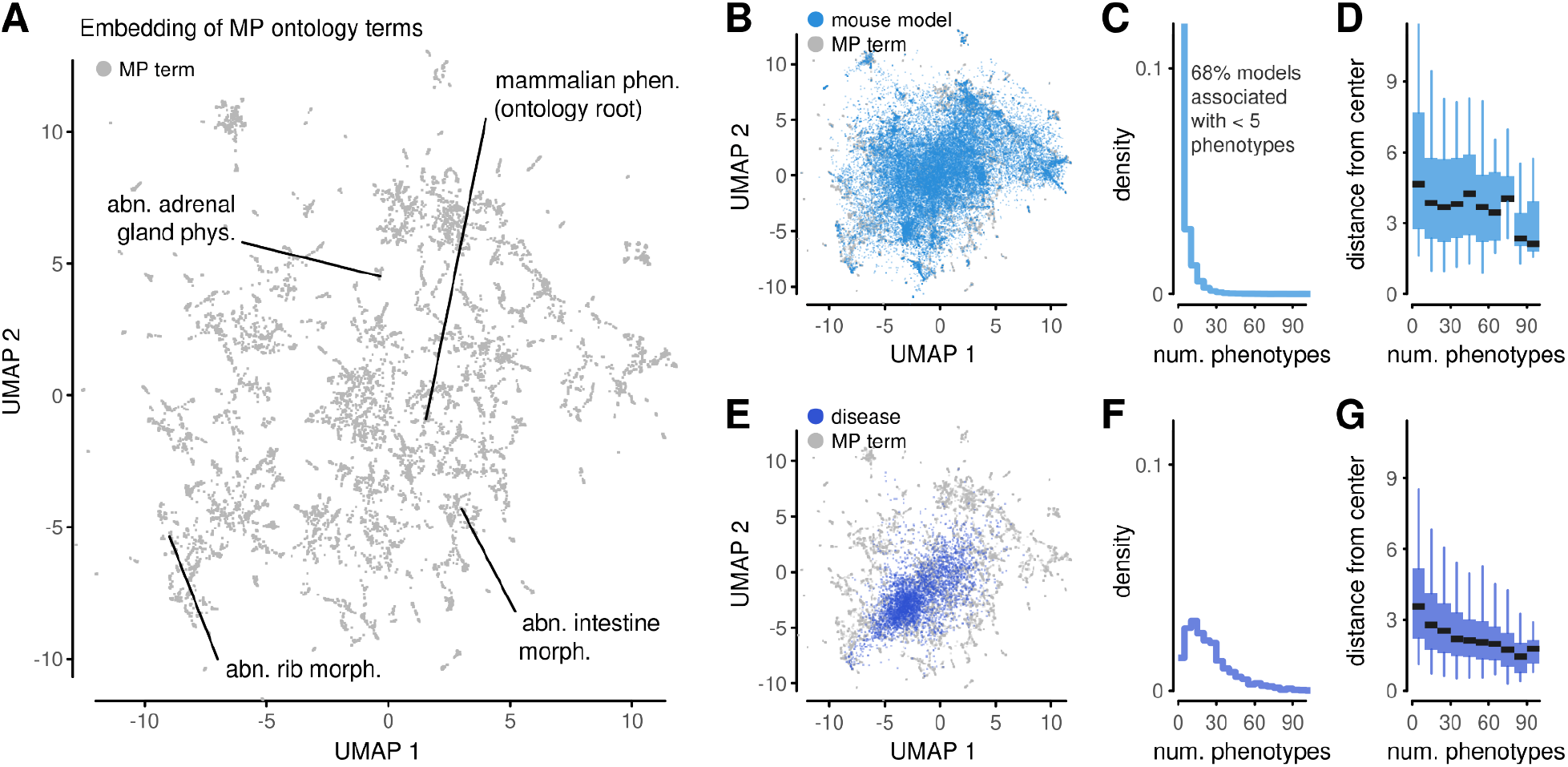
Embeddings of mammalian phenotypes. (A) Embedding of mammalian phenotype (MP) ontology terms based on text similarity. Labels point to selected ontology terms. (phen.: phenotype, abn.: abnormal, dev.: development) (B) Projection of mouse models into an embedding of ontology terms via averaging of coordinates of their annotated phenotypes. (C) Histogram of the number of phenotypes for all mouse models. (D) A summary of the position of mouse models in the projections in (B), stratified by the number of annotated phenotypes. Boxes represent 25%-75% intervals, whiskers represent 5%-95% intervals, middle lines represent medians. (E-G) Analogous to (B-D) using phenotype profiles of human diseases translated into the mammalian phenotype ontology.

We assembled annotations about animal models from the Mouse Genome Database (MGD) (Blake et al., 2021) and the International Mouse Phenotyping Consortium (IMPC) (Dickinson et al., 2016). This resulted in a collection of 53,629 models that describe mouse strains with mutations in one of 15,729 distinct genomic markers (Methods). Most markers correspond to protein-coding genes or non-coding genes, but also include other genomic constructs; we treated all on an equal level. The models carried 254,623 annotations to 9,907 different MP terms. We computed the position of these models in the embedding space by averaging the coordinates associated with their phenotypes. These projections appeared throughout the embedding space (Figure 1B), conveying that the animal models have diverse phenotypic features. Next, we investigated the distribution of the number of phenotypes per model. 86% of models were associated with fewer than ten phenotypes and 69% with fewer than five (Figure 1C). Despite the large skew, 31,477 models were associated with between two and 113 MP terms, allowing us to stratify the dataset. The number of annotations created a bias in the position of the models within the embedding: the distance of the model from the center was anti-correlated to the number of phenotypes (Figure 1D). This is an indication that of the two coordinates in the visualization, one is effectively taken up to capture the number of annotations rather than their biological meaning.

We performed analogous calculations for phenotype profiles of human diseases (Methods). We translated disease phenotypes into sets of MP terms, and then projected the translations into the embedding space (Figure 1E). Diseases were, on average, linked with more phenotypes than mouse models (Figure 1F) and also exhibited a correlation between the number of phenotypes and the distance from the center (Figure 1G). Indeed, the bias was more marked than for mouse models. The bias creates an impression that diseases with rich annotations are more similar to one another than diseases with few annotations, which is not justified from a phenotypic perspective. It also drives the distribution of profiles into a unimodular shape, which does not capture the diversity and multi-dimensionality of human diseases.

While projecting phenotype sets into the ontology embedding through coordinate averaging may produce insight for small phenotype sets, our results demonstrate that this approach tends to place well-annotated phenotypic profiles toward the center. The bias becomes more evident as the number of annotations increases; projections of diseases are more affected than mouse models. As the bias is related to averaging, it is bound to appear with other embeddings of MP terms as well, for example those generated based on the ontology hierarchy graph. Furthermore, because phenotypic annotations are expected to become more detailed with time, such bias should be expected to grow as well. Thus, embeddings of ontology terms in low dimensions are not recommended for the exploration of complex phenotype profiles.

### Embeddings of full phenotype profiles capture the diversity of animal phenotypes

As an alternative to treating mouse models as sets of phenotypes and averaging coordinates for ontology terms, we produced embeddings for the mouse phenotype profiles directly. There are several possible approaches to encode phenotype profiles into a numerical form that can be processed with dimensional reduction algorithms (Figure S2).

In a first attempt, we constructed vector representations for individual mouse models using a previously described procedure (Konopka & Smedley, 2020). This approach estimated prior probabilities for all phenotypes based on their frequency in the cohort, and then integrated annotations from model-specific phenotypes to create real-valued vectors. This approach thus combined information from the entire cohort, from model-specific annotations, and from the ontology hierarchy. We applied UMAP for dimensional reduction and summarized 53,629 mouse models from MGD and IMPC into a complex layout (Figure 2A). The embedding was not dominated by technical variables such as data source, mouse genetic background, or zygosity of genetic mutations (Figure S3). It was, however, influenced by the number of model phenotypes (Figure S3). It separated some models with few phenotypes from models with rich annotations, but also created a complex layout among models with many phenotypes. Overall, the embedding thus provides a concise visualization of the phenotypic diversity in the dataset.

**Figure 2.**
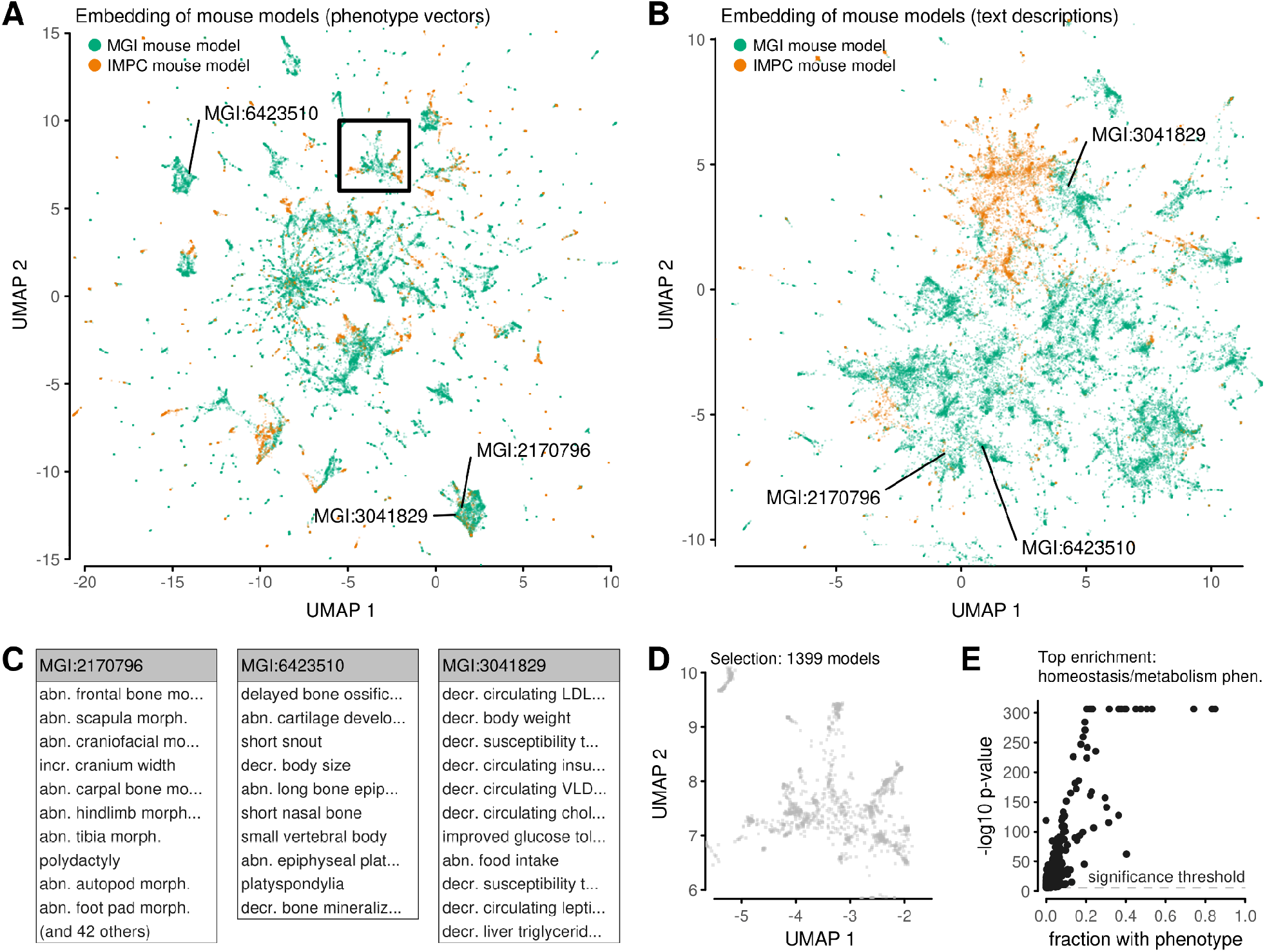
Embeddings of mouse models. (A) Embedding of mouse models based on vector representations of their phenotypes. Models are colored by the source of curated data. Labels and the rectangle point to selected models. (B) Analogous to (A), but with the layout based on semantic similarities of text descriptions. (C) Lists of phenotypes associated with individual mouse models highlighted in (A). Some lists are truncated for this visualization. All phenotype names match definitions from the ontology (abn.: abnormal, morph.: morphology, incr.: increased, decr.: decreased). (D) A magnification of a small region of the embedding in (A). (E) Enrichment analysis comparing the phenotypes associated with mouse models in (D) against models outside of the selected region. Dots correspond to phenotypes in the ontology. Statistical significance (p-value) is evaluated using the Fisher test; the significance threshold is Bonferroni-corrected p=0.05. The most significant phenotype is labeled.

For comparison, we produced embeddings with alternative approaches spanning several mathematical techniques: using binary phenotype vectors, text-based semantic similarities, and graphs (Methods). These approaches all used more coarse-grained representations of phenotypes than our real-valued vectors. They resulted in embeddings that were all substantially different from one another (Figure S2). In particular, graph-based methods produced homogeneous layouts that did not capture any patterns among mouse models, so they were not considered further.

One of the approaches that emphasized differences between groups of models used text-based semantic similarities (Figure 2B). Similarly to the first embedding, this approach suggested models are part of groups of various sizes (Figure 2B). However, relative distances between mouse models differed: models that appeared nearby in the first embedding were far-apart in the second, and vice versa (Figure 2B). Such disparities are not unexpected given the ambiguities in turning phenotype annotations into a numerical representation. The text-based embedding also showed a more prominent separation of models according to MGD or IMPC data source. This may arise because IMPC data, which originate from a systematic screen rather than from bespoke experiments, have a more limited range of MP terms than MGD models. These MP terms may have influenced text-based calculations, which detect similar phrases in model descriptions, more than vector based calculations, which utilize the ontology hierarchy in a more direct manner.

Each embedding is rich in interpretable information. At a fine-grained level, each data point corresponds to a specific phenotype profile (Figure 2C) (Lyon et al., 1996; Hayes et al., 1998; Schreyer et al., 2001; Grigelioniene et al., 2019). Furthermore, each embedding is interpretable at a regional level. To demonstrate such patterns, we selected a region in our first embedding (Figure 2D). Enrichment analysis revealed an over-representation of certain phenotypes among the models within that region (Figure 2E). Such enrichment is not surprising as the embedding was constructed so that similar phenotypes appear together. Indeed, other examples also exhibited enrichment (Figure S4), validating the embedding as well as providing a mechanism to assign regions with phenotypic interpretations.

These experiments demonstrate that our calculations produced more than one reasonable low-dimensional embedding of mouse models. Although it is unclear how to choose a single embedding as a reference map for mouse phenomics, these candidates can be used for data exploration.

### Neighborhoods provide insight for phenotype prediction

Having demonstrated that embeddings summarize the diversity among mouse models and reveal qualitative patterns, we next investigated whether they can provide quantitative insight for individual models. To this end, we computed predictions of phenotype profiles based on nearest neighbors (Figure 3A). For each model, we extracted a ranked list of nearest neighbors in the high-dimensional representations and in low-dimensional embeddings. We averaged the vector representations of the neighbors to create a prediction, and evaluated the error between the prediction and the model’s true representation (Methods). We then investigated the mean prediction error as a function of the number of neighbors, denoted as k (Figure 3B). This approach is used in self-supervised learning to calibrate free parameters without a ground-truth (Batson & Royer, 2019). For predictions based on neighbors evaluated from the original data, i.e. from high-dimensional vectors, the optimal k was k=2. For approaches based on embeddings, the optimal number of neighbors varied with the embedding dimension, but all were below k=10. As expected, prediction errors were lowest when the neighbors were computed from the high-dimensional data, and highest when the neighbors originated from two-dimensional embeddings used for visualization.

**Figure 3.**
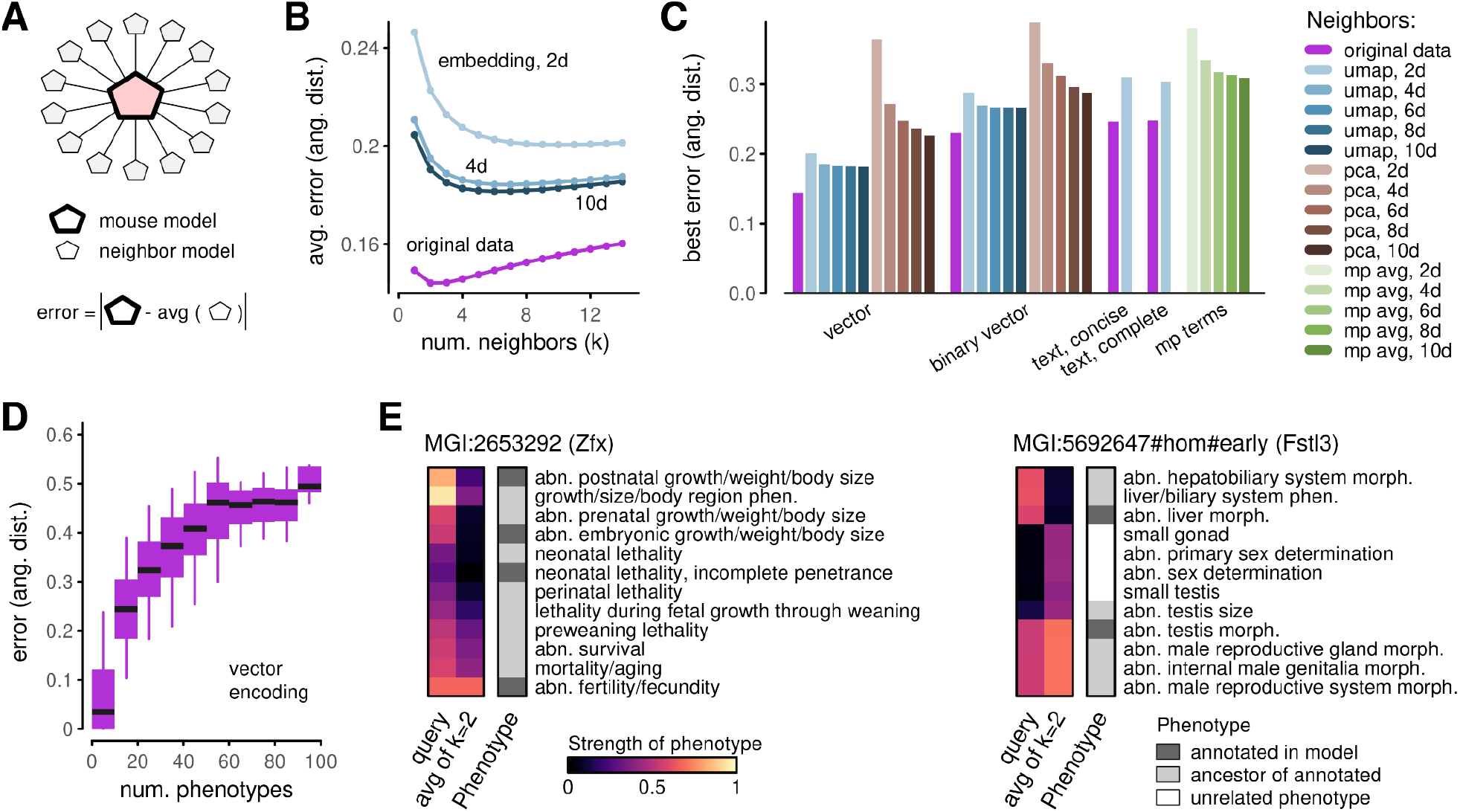
Phenotype prediction. (A) Schematic explaining phenotype prediction using neighbors. Given a mouse model, its predicted phenotype profile is defined as a simple average over its neighbors. An error is defined as the L2 norm between the model profile and the prediction. (B) Exploration of mean prediction error as a function of the number of neighbors used in the calculation. Lines correspond to distinct ways of identifying neighbors: from original vector representations, or from embeddings in various dimensions. (C) Summary of best-achieved errors for prediction approaches using original vector data, original binary vector data, embeddings in various dimensions, and using text-based similarity measures. (D) Stratification of mouse models by the number of model phenotypes. Boxes represent 25%-75% intervals, whiskers represent 5%-95% intervals, middle lines represent medians. (E) Examples of mouse model phenotype vectors and predictions based on two nearest neighbors. Heatmaps only show a small number of phenotypes that contribute the most to prediction errors. Categorical phenotype annotations indicate whether a listed phenotype is one of the models’ annotated phenotypes, an ancestor of an annotated phenotype, or a phenotype unrelated to model annotations.

To further investigate the factors that can affect the prediction of phenotypes, we repeated these calculations with a series of approaches. We computed neighbors using high-dimensional representations based on real-valued (non-binary) phenotype vectors, binary phenotype vectors, and text descriptions. We considered dimensional reduction via UMAP, principal component analysis (PCA), and coordinate averaging of MP terms. For each combination of data encoding and dimensional reduction technique, we calibrated the optimal number of neighbors and reported the mean prediction error (Figure 3C). All errors were computed with respect to the non-binary vector representations. Given this choice, it is unsurprising that the smallest prediction error was achieved by the approach that used neighbors from high-dimensional non-binary vector data. However, this choice is useful because it ensures that the scales for all errors are comparable.

Among approaches that used dimensional reduction, we observed an improvement (reduction) in prediction error with increasing embedding dimension, d, for all dimensional reduction techniques (UMAP, PCA, MP coordinates). Interestingly, the improvement plateaued quickly with UMAP; there was little change between d=8 and d=10 dimensions. In contrast, prediction errors from PCA showed a steadier improvement with d. However, errors from PCA embeddings were higher than for UMAP, and remained higher for PCA with d=10 than for UMAP with d=2.

The encoding scheme (non-binary phenotype vectors, binary phenotype vectors, text descriptions) had a substantial effect on prediction errors. Variability between encodings was large compared to differences between UMAP in various dimensions. This indicates that studies of mouse models are bound to be more influenced by how phenotype data are curated and encoded than by any loss of resolution due to dimensional reduction. At the same time, prediction errors were relatively similar for binary vector-based and text-based approaches.

Average prediction errors hide variation within the cohort. To investigate heterogeneity in more depth, we stratified models according to the number of associated phenotypes (Figure 3D). Errors for models with few phenotypes were generally low. This is because when a model has a limited number of phenotype annotations, there may exist other models with the same characteristics. As an example, our dataset had 14 models annotated with the single phenotype of “deafness”. Averaging a small number of nearest neighbors that may have equivalent phenotypes produces predictions with zero error. For models with more annotations, the probability that other models have the same set of phenotypes by chance decreases.

Predictions that deviate from the annotated model representation highlight which of a model’s phenotypes are unusual. They can also suggest phenotypes that may be missing in the annotations. To illustrate such reasoning, we visualized the discrepancies between two models and their predictions (Figure 3E) (Luoh et al., 1997; Dickinson et al., 2016). In each case, part of the discrepancies originate from different weights assigned to measured MP terms or their ancestors, i.e. MP terms that are upstream in the ontology hierarchy. Other discrepancies arise from phenotypes that are not related to the models’ annotations. If nearest-neighbors were to be used as a denoising scheme, these discrepancies would result in imputed phenotypes. Even without formal imputation, if embedding coordinates were used as inputs for a machine-learning classifier, these phenotypes would influence how the model would be utilized by the classifier. Thus, the neighbor prediction can be informative for interpreting, or explaining, outcomes of downstream ML models.

### Neighborhoods highlight consistency as well as heterogeneity of genotype-phenotype annotations

Models in our dataset are characterized by the mouse background strain, mutation of a single genetic marker, and mutation zygosity. While combinations of these features appear uniquely in the dataset, several models can be linked to the same genetic marker. A summary of the number of models per marker, which we refer to as genes for simplicity, revealed a skewed distribution (Figure 4A). Thousands of genes were represented by a single mouse model in the dataset, but thousands of others were mutated more than once. The three most studied genetic markers appeared in more than 200 models each (Figure 4A, inset).

**Figure 4.**
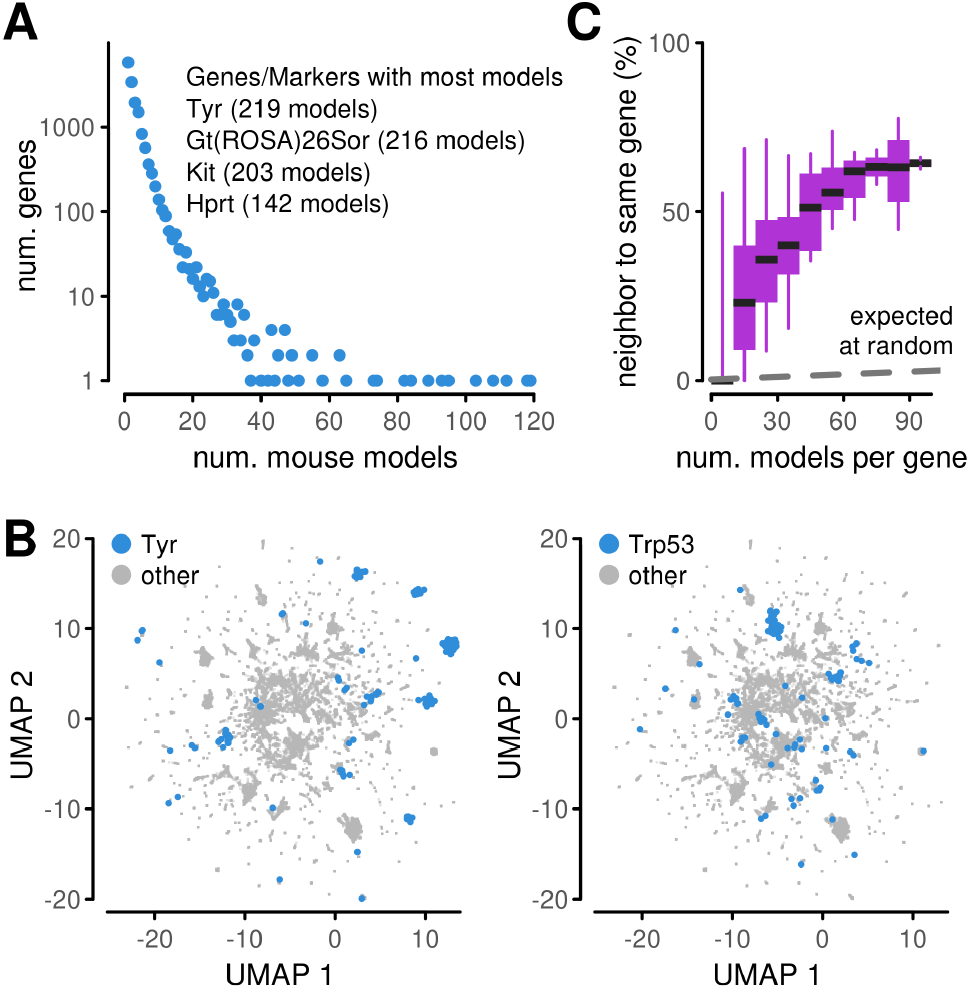
Phenotype heterogeneity. (A) Multiplicity of models available for individual genes. The genes represented in the most models are listed in the inset. (B) Embeddings of mouse models highlighting the location of models with selected genes knocked-out. Highlighted models are jittered to better display the number of models. (C) Proportion of genes for which the nearest-neighbors of a mouse model contain another model with the same gene knocked-out. The summary is stratified by the number of models available for a gene. Boxes represent 25%-75% intervals, whiskers represent 5%-95% intervals, middle lines represent medians. Dashed line indicates an expected level under a null hypothesis that neighbors are selected at random.

The multiplicity of models linked with the same gene provides opportunities to study consistency and heterogeneity among genotype-phenotype associations. For illustration, we picked some of the most studied genes and highlighted their models in embeddings (Figure 4B, Figure S5). Some of the models appeared close together. This suggests consistency in the phenotypic annotations linked to the gene and robustness with respect, for example, to mutation construct. At the same time, models linked to one gene also split into several close-knit clusters in distinct parts of the embedding. This suggests an opposite effect, i.e. heterogeneity in annotations. Heterogeneity may be due to incompatible curation, incomplete phenotyping measurements on some of the models, or differences in phenotype due to the genetic background (Figure S5). The embedding cannot deconvolute these effects, but the visualization provides qualitative insight on the scale of the phenomenon in the mouse data.

To quantify the extent of consistency and heterogeneity, we studied the nearest neighbors of all models and assessed the proportion of times a model was near another model with the same gene (Figure 4C). This fraction increased with the multiplicity of mouse models. Among genes represented by more than 20 models, the proportion of models linked to another model with the same gene was above a third. This was substantially higher than expected if neighbors were assigned by chance. Nonetheless, since the level was below 0.5 for most genes, it is more likely for a randomly selected model to have neighbors with different genetic composition than to link to at least one model with the same mutated gene. This can be due to difficulties in encoding the data, incompleteness in the dataset, or due to overlapping phenotypes associated with different genes.

### Embedding diseases alongside animal models provides grounds for exploring disease-causing genes

Finally, we returned to the dataset of human diseases. For those diseases with annotations in the form of HP terms, we translated the phenotypes into MP terms and constructed vectors-based and text-based representations. We then projected the diseases into our previously-generated embeddings (Methods). With both vector-based (Figure 5A) and text-based (Figure 5B) approaches, diseases covered large areas of the embedding space and formed several disjoint clusters. Patterns were robust to how the disease phenotypes were translated from human to mammalian phenotype ontologies (Figure S6). Interestingly, disease projections avoided certain areas of the embeddings. For example, one of the areas omitted by the diseases was enriched in phenotypes related to mortality and ageing (Figure S4). Diseases linked to well-annotated models with all genetic backgrounds and showed a slight preference toward models with homozygous mutations (Figure S6).

**Figure 5.**
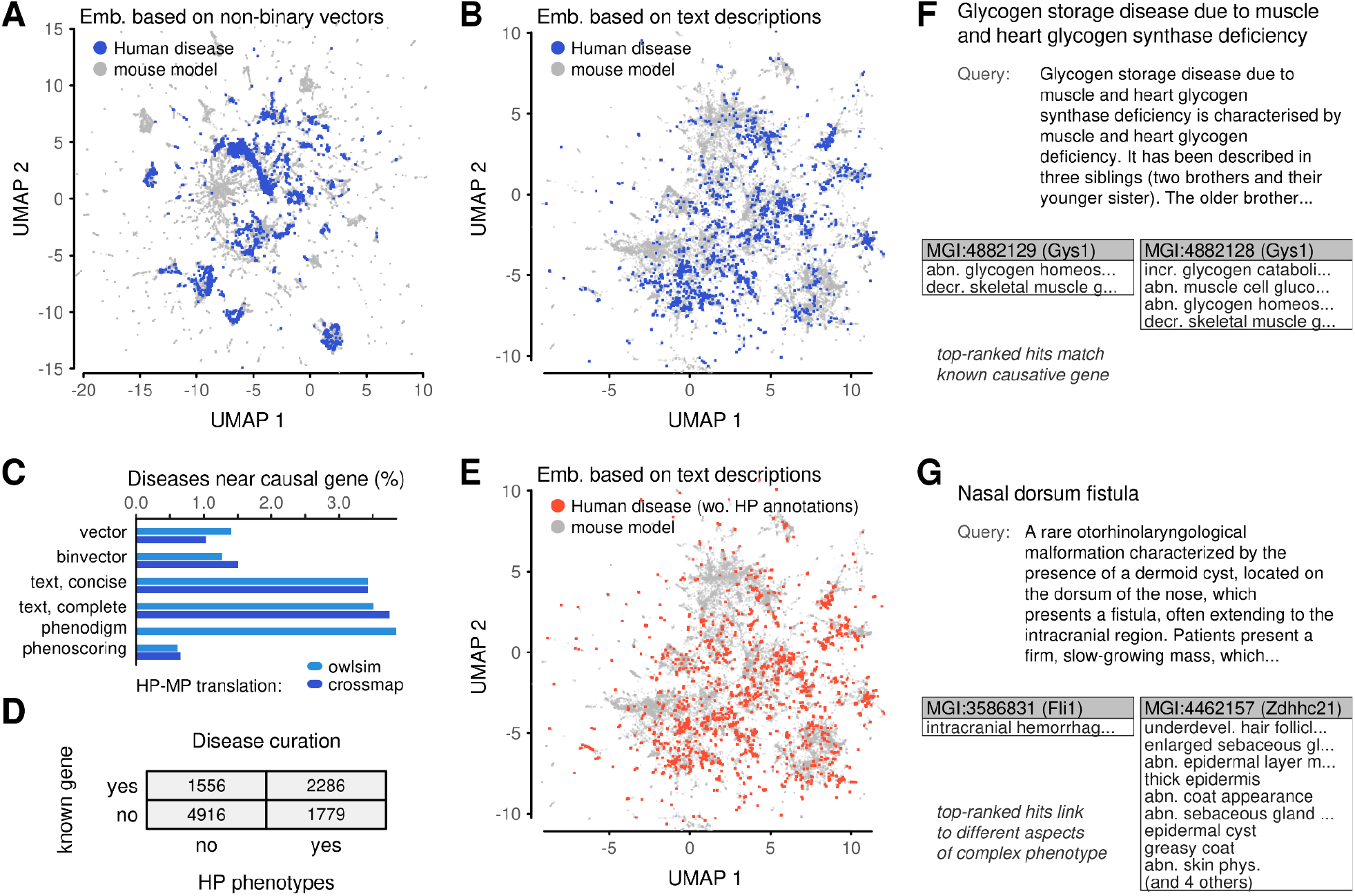
Embedding of human diseases in the mouse phenotypic space. (A) Projection of human diseases into an embedding of mouse models based on phenotype vectors. (B) Analogous to (A), but with the underlying embedding produced based on semantic similarities of text-based descriptions. (C) Summary of causal-gene extraction. Diseases with phenotype and gene annotations were compared with all mouse models. The percentage in the bar graph is the proportion of diseases for which one of the k=15 nearest mouse models contained a mutation in the causal gene. (D) Summary of ORPHANET disease annotations in terms of phenotype ontology terms and causative genes. (E) Projection of human diseases without HP annotation into an embedding of mouse models based on text similarity. (F,G) Examples of text-based disease descriptions along with two mouse models, selected manually from among the top five search hits.

Next, we asked to what extent similarities between diseases and mouse models can link diseases to their associated genes. For simplicity, we called all disease-associated genes as causative genes. We compared approaches based on numeric vectors, based on text descriptions, and two previously described methods for scoring disease-model associations (Smedley et al., 2013; Konopka & Smedley, 2020). The proportion of diseases that had a mouse model with the causative gene within 15 nearest neighbors was low: below ∼4% (Figure 5C). Interestingly, the algorithm used to translate between human and mammalian phenotype ontologies had a smaller effect than the data encoding or the algorithm for computing neighbors.

The best performer was a scheme specifically designed to score disease-model associations (Smedley et al., 2013). Strikingly, text-based approaches performed almost on par, better than vector-based approaches.

Given that text-based descriptions performed well in matching diseases to causative genes, we reasoned that this approach could be a gateway to studying diseases without formal phenotype annotations. Among the dataset of human diseases, 61% did not have any HP annotations (Figure 5D). These diseases could not be included in calculations that rely on ontologies. They were also excluded from the gene-mapping assessments to ensure like-for-like comparisons (Figure 5C). However, these diseases have text descriptions, so they can be used in text-based calculations. Projections into the embedding of mouse models revealed they fell into the same regions as before (Figure 5E). This indicates that un-annotated diseases span the whole phenotypic space, and also that our treatment of text descriptions did not introduce excessive biases that would place these cases apart from well-annotated diseases. In more depth, the un-annotated diseases linked to mouse models of all genetic backgrounds and annotation levels, albeit with a preference to mouse models with few phenotypes (Figure S6).

To illustrate explorative analyses based on text descriptions, we searched for two diseases without formal phenotype annotations (Figure 5F, 5G). An initial search for a glycogen deficiency (ORPHA:137625) correctly linked this disease to mouse models with abnormalities in glycogen homeostasis (Figure 5F). Because of the naive treatment of text in our calculations, some of the top-ranked models were characterized by opposite directional effects to the disease description. Among the top hits were models with mutations in *Gys1* (Bouskila et al., 2010), the known causative gene for the disease. Other hits with similar phenotype profiles had mutations in *Ppp1r3a* and *Gyg*, both genes that participate in glycogen homeostasis. This confirms that text search can link human diseases to relevant mouse data. In this case, the top search hits all had consistent phenotype profiles, so a projection of the diseases into low-dimensional embeddings can also be expected to link the disease with a neighborhood of relevant mouse models.

Separately, we searched for an otorhinolaryngological disorder (ORPHA:141219) characterized by cysts around the nose and extending into the cranium (Figure 5G). The causative gene for the disorder is not known. The first two hits in text-based search matched the disease to mouse models with quite distinct phenotypes. The first was characterized by an intracranial phenotype (Hart et al., 2000); the second by phenotypes of the epidermis (Mill et al., 2009). Such hits can appear in distinct portions of an embedding. Projecting the diseases into the space of models might place the disease close to one at the expense of the other. This can produce discrepancies between sets of neighbors computed in the original high-dimensional space and the low-dimensional embedding. This is a documented effect inherent to dimensional reduction (Cooley et al., 2020). In the disease context, it highlights the value in scrutinizing raw search results in addition to a low-dimensional visualization.

## Discussion

Animal models offer a direct route to characterizing the impact of genetic mutations. While studying the relationship between genotype and phenotype is often performed gene-by-gene, careful curation of the literature (Blake et al., 2021) as well as systematic phenotyping of hitherto-unstudied genes (Dickinson et al., 2016) mean that the collective data for the mouse will approach whole-genome coverage in the near future. This opens possibilities to utilize mouse phenotypes as a reference dataset in genomic analyses. As such, it is important to characterize the potential such a dataset offers for data exploration, machine learning, and downstream applications. In this work, we explored dimensional reduction for this data. The results visualize the heterogeneous landscape of mouse phenotypes. Our calculations also provide qualitative and quantitative observations about the strengths and limitations of this pool of data.

A challenge in dealing with large-scale phenotype data is that there are several plausible ways to encode sets of phenotypes so that they can be used for calculations. We explored vector-based, text-based, and graph-based approaches. There also exist many algorithms that can perform dimensional reduction. We focused on approaches that can be used for visualization and thus focused on dimensional reduction into two dimensions. Such embeddings enable interactive, human-led data exploration. However, we also investigated embeddings into higher dimensions, which can be beneficial for machine learning.

Strikingly, certain strategies provide sub-optimal visualizations in two dimensions. A strategy that first creates an embedding of an ontology and then projects phenotype sets into that space via coordinate averaging is prone to construct visualizations that are dominated by technical features, notably the number of phenotypes within a phenotype set (model or disease). This result has a mathematical justification: averaging summarizes heterogeneous elements, in this case phenotypes, to a central value, with the variability of the outcome decreasing with the number of elements. It is also worth comparing this strategy with analysis pipelines in transcriptomics, which do not create embeddings based on genes and project expression profiles for samples into that space, but rather create embeddings for samples directly. Another suboptimal strategy is one that uses the MP ontology as a graph and performs a joint embedding of the ontology and a set of mouse models. This approach hides much of the rich phenotypic similarities and differences among models (Figure S2).

Strategies based on phenotype vectors or text descriptions produce visualizations that are interpretable at the level of single mouse models and group similar models together. These visualizations can thus be said to capture the phenotypic diversity among mouse models in the available data. However, they can also reflect peculiarities and limitations of the underlying data. Embeddings show many small, isolated groups with peculiar combinations of MP terms. Such combinations may arise due to targeted phenotyping efforts on those mouse models rather than true phenomenological specificity compared to other mutants. Small annotation sets may also arise when phenotyping is carried out as part of a high-throughput screen (Dickinson et al., 2016). Such annotation sets may grow in time as further observations are catalogued and curated from literature (Blake et al., 2021). Analysis methods can be designed specifically to track changes of annotations due to steady growth (Konopka & Smedley, 2020). However, in the context of dimensional reduction, changes in the annotation set should be expected to disrupt the positioning of individual models within an embedding. Thus, the embeddings should be expected to be unstable for models with low annotation counts. The overall structure of the embeddings may change too, albeit to a lesser extent.

Besides providing visualizations of the heterogeneous phenotype data, we investigated schemes for phenotype prediction based on nearest neighbors. These predictions are informative from at least three perspectives. First, they suggest new experimental assays on individual mouse models that may complete their phenotype profiles. Second, the predictions can be used for interpreting outputs from machine learning models trained using embedding coordinates. Third, we used prediction errors to quantify the information loss produced by various data-encoding and data-embedding approaches. The results confirm expected properties, namely that embedding data into low-dimensional spaces loses some information, that increasing the target dimension increases the fidelity of the embeddings, and that non-linear methods like UMAP preserve more information than linear methods like PCA. Interestingly, the results also show that discrepancies between predictions from original data and from embeddings can be comparable to discrepancies between different encodings of the original data (e.g. binary or real-valued vectors). This suggests that any analyses based on phenotype data are likely to be sensitive to how the raw data is prepared. Indeed, they may be more sensitive to data preparation than to dimensional reduction.

We corroborated this sensitivity in calculations projecting human diseases into embeddings of mouse models. We used nearest neighbors to link human diseases to mouse models and thereby to genes. Recall of established disease-gene associations varied depending on the encoding strategy (vector, text, etc.). Interestingly, approaches based on text similarities were among the most performant. Considering that these approaches are tunable (Konopka et al., 2021), can integrate datasets other than phenotypic annotations, and that they execute two orders of magnitude faster than a dedicated scoring scheme for scoring disease-gene associations, they represent promising avenues for subsequent analyses.

## Methods

### Phenotype data

Definitions of the human phenotype (HP) ontology (Köhler et al., 2018) and the mammalian phenotype (MP) ontology (Smith & Eppig, 2015) were obtained through the OBO Foundry (HP version 2021-02-28, MP version 2021-01-12). Terms from the HP ontology were mapped onto terms in the MP ontology using owlsim (Washington et al., 2009), which is an ontology-aware algorithm, and using crossmap (Konopka et al., 2021), which performs searches based on text similarity. Mappings with crossmap were performed using diffusion driven by the MP dataset and by a set of manual annotations (Konopka et al., 2021). Both owlsim and crossmap map queries to multiple hits. Translations between the HP and MP ontologies were established using only the best-ranked mapping.

Mouse model definitions and associated phenotypes were obtained through the data portal of the International Mouse Phenotyping Consortium (data release 14.0) (Dickinson et al., 2016). Data downloaded from the IMPC included definitions of mouse models curated by the Mouse Genomics Database (Blake et al., 2021). The dataset contained information about lines with mutations in only one marker or gene each; the dataset did not cover mouse models with extended mutations affecting multiple genes.

Disease definitions and associated phenotypes were downloaded from Orphadata (http://www.orphadata.org; data version 2021-04-01) and parsed using custom scripts (https://github.com/tkonopka/crossmap).

### Ontology representations and embeddings

Embeddings for ontology terms were generated using two approaches (Figure S1). In a first approach, text strings were constructed for each term in the MP ontology by concatenating phenotype name, definition, synonyms, and comments. These strings were loaded into a crossmap knowledge-base (Konopka et al, 2021; https://github.com/tkonopka/crossmap). The crossmap instance splits text into bags of k-mers, weights the k-mers according to their information content, and builds a nearest-neighbors index. The crossmap instance was used to perform searches and compute sets of nearest neighbors for each ontology term. The nearest neighbors were provided to the UMAP algorithm (Becht et al., 2018) implemented in R to produce embeddings in two dimensions. Settings were left at the default values, except for knn_repeats=3 to increase the quality of nearest-neighbor search, and min_dist=0.2 to increase space between adjacent points (for visualization).

In a second approach, MP ontology terms were treated as nodes in a graph. Edges between nodes were set if two MP terms were linked by ‘is a’ relationships in the ontology hierarchy (which is the only relationship type defined in the MP ontology). The resulting graph was processed using node2vec (Grover & Leskovec, 2016) to produce embeddings in two dimensions. Embeddings were produced using the snap implementation (https://github.com/snap-stanford/snap) with default parameters and the python implementation (https://pypi.org/project/node2vec/) with default settings. The python implementation was also executed with non-default settings with num_walks=5 and walk_length=5.

Both UMAP and node2vec are stochastic algorithms and embeddings may differ when repeated. All calculations were performed twice with two different seeds for random number generation. Consistency between replicates and between different embedding approaches was assessed by extracting sets of 15 nearest neighbors in the low-dimensional embeddings, and computing the mean Jaccard indexes.

### Mouse model representations

Mouse models were defined as sets of phenotypes associated with a specific mouse strain (genetic background), a single-gene knock-out mutation, and zygosity. For IMPC data, models were further subcategorized by life stage (embryonic, early-adult, or late-adult).

Raw data for each mouse model were encoded in several ways: based on numeric vectors, on text, and using graphs (Figure S2). The numeric representations consisted of vectors of length equal to the size of the MP ontology. In a binary approach, values were set to zero by default and changed to one if an MP term was linked with a mouse model, or was an ancestor of such an MP term. Non-binary vector representations were constructed following a published protocol (Konopka & Smedley, 2020). Briefly, values within the vectors were initially set to prior probabilities for each MP phenotype, which were estimated from the ensemble of non-IMPC models. Values were updated through a Bayesian procedure with phenotype annotations and then propagated using the ontology hierarchy.

Two types of representations were constructed starting from text. A ‘concise’ representation was defined by concatenating the names of all MP terms associated with a model. A second ‘complete’ representation was constructed in the same way, but also including the names of all ancestors of MP terms associated with a model. Text strings defined in these ways were loaded into an instance of a crossmap knowledge-base (Konopka et al., 2021). This software uses k-merization to turn text into numeric representations. Weights for k-mers were computing using a corpus of text including dictionary definitions of English words (www.wiktionary.org) and phenotype definitions from the MP ontology.

For graph-based approaches, MP terms and mouse models were used as graph nodes. Edges were defined between MP nodes if the corresponding MP terms were linked by ‘is a’ relationships in the ontology. Edges were defined between mouse nodes and MP nodes from mouse model associations.

### Mouse model embeddings

Embeddings of mouse models based on vector and text representations were performed with the UMAP algorithm (Becht et al., 2019) through an R implementation.

For encodings based on non-binary and binary vectors, all vectors were normalized to unit norm and then provided to the embedding algorithm. Settings were left at default, except for knn_repeats=3 to increase the quality of nearest-neighbors and min_dist=0.2 to spread points for visualization purposes. In calculations exploring the impact of the embedding dimension, UMAP runs were provided with the same set of nearest neighbors, thus eliminating stochasticity effects due to approximate calculations of neighbors.

For encodings based on text, raw data were managed with a crossmap instance (https://github.com/tkonopka/crossmap). The crossmap instance was used to search for sets of nearest neighbors. Nearest neighbors were provided to the UMAP algorithm in R, which was run as described for the vector approaches. For graph-based approaches, embeddings were generated using node2vec (Grover & Leskovec, 2016) through the snap implementation (https://github.com/snap-stanford/snap) and python implementations (https://pypi.org/project/node2vec/) with default settings. The python implementation was also run using non-default settings walk_length=5 and num_walks=5. The graph-based approaches produced a joint embedding of MP terms and mouse models; only mouse models were used in visualizations.

Calculations of nearest neighbors with crossmap and embeddings with UMAP and node2vec rely on stochastic algorithms. All calculations were performed with two different seeds for random number generation. Consistency between replicates and between different approaches were assessed as for embeddings of ontology terms. Sets of 15 nearest neighbors were computed in the low-dimensional spaces, and Jaccard indexes were computed for matched points in the various embeddings.

While all embeddings were computed based on all the available data, some visualizations were truncated to enhance the presentation. Models that were not associated with any phenotype (MP:0002169, ‘no abnormal phenotype detected’) were excluded from all visualizations, as were models with only one phenotype. Visualizations excluded these models to focus attention on 31,477 models with rich annotations. (The exception is Figure S3, which includes both zero-phenotype and one-phenotype models.) Visualizations also set quantile-based limits for axes to focus attention on the central parts of embeddings with. Quantile intervals were at least as wide as 2%-98% for each axis.

### Projections of human diseases into embeddings of mouse models

Human diseases with phenotype annotations from the HP ontology were translated to the MP ontology by replacing their HP terms with best-matching MP terms. Sets of MP terms derived from diseases were encoded into non-binary vectors using the same procedures as for mouse models. Vectors with disease profiles were compared with all mouse models to identify nearest neighbors, and the position of each disease in an embedding was computed using UMAP. This calculation initially places a disease at the averaged location of its nearest mouse models, and subsequently adjusts this position using the UMAP optimization algorithm.

For encodings based on text, human diseases were taken to consist of a clinical summary paragraph, the names of associated phenotypes (MP translations), and the names of curated genes. These text documents were compared with phenotype descriptions of mouse models using crossmap to produce sets of nearest mouse models. Projections of the disease descriptions in embeddings of mouse models were defined as the coordinate of the single most-similar mouse model. A more sophisticated approach utilizing several mouse models was not possible in this case due to a technical limitation of the R UMAP package.

## Software and Data Availability

Source code for analysis scripts is available on GitHub at https://github.com/tkonopka/mouse-embeddings. The data underlying this article are available in Zenodo at https://doi.org/10.5281/zenodo.4916171.

### Funding

This work was supported by a grant from the National Institutes of Health [Grant 5-UM1-HG006370 to D.S.].

## Supplementary Figures

**Figure S1.**
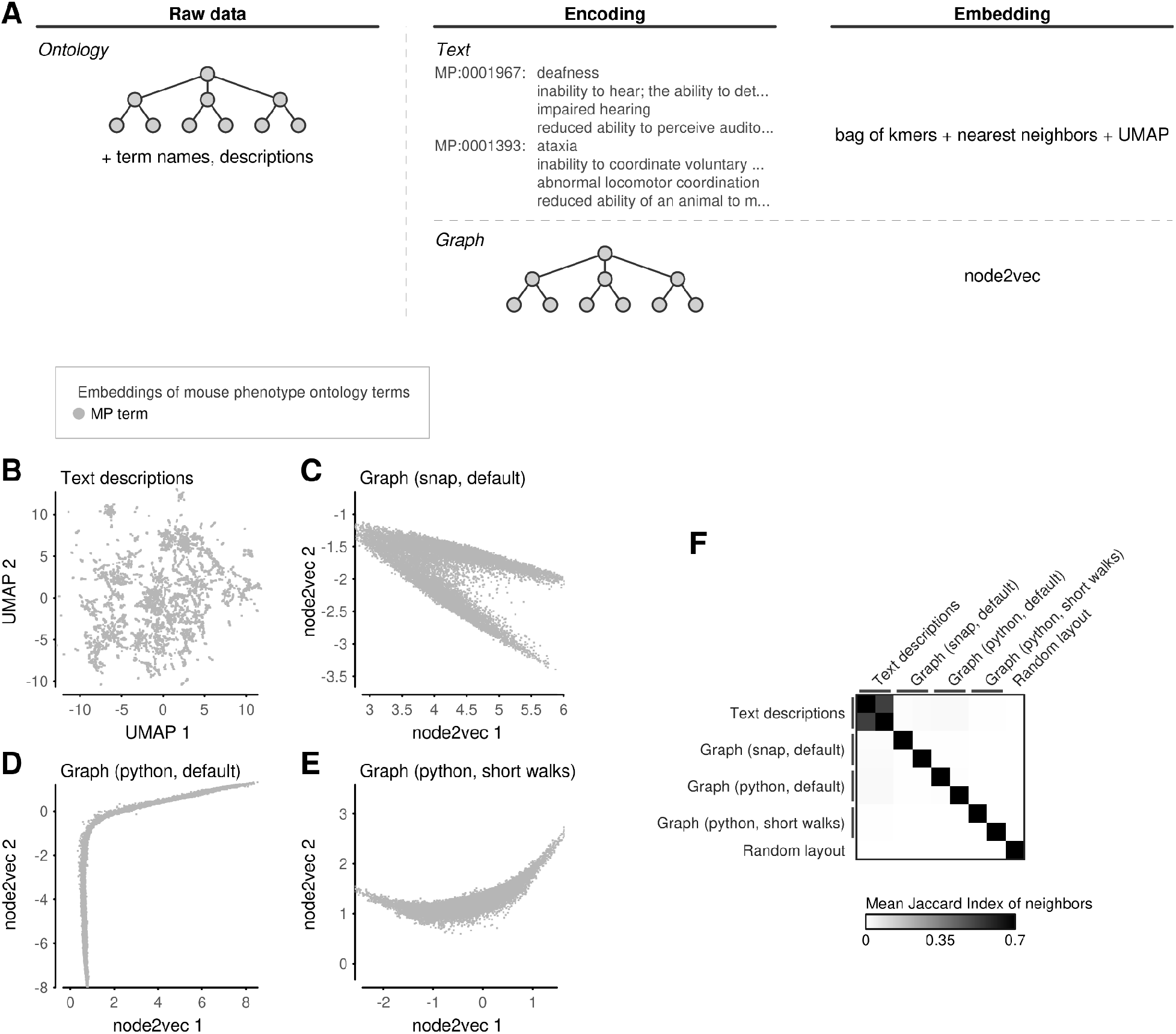
Embeddings of mammalian phenotype ontology terms. (A) Schematic of ontology data, possibilities for data encodings, and algorithms for creating embeddings. (B) Embedding of mammalian phenotype (MP) ontology terms based on text descriptions (name, definition, synonyms, comments, and name of parent term). Similarities computed using crossmap and layout generated with UMAP. (C) Embedding based on the hierarchy relations between ontology terms, generated using snap implementation of node2vec with default settings. (D) Embedding based on the ontology hierarchy graph, generated using python implementation of node2vec with default settings. (E) Similar to (D), generated with settings walk-length and num-walks set to 5. (F) Comparison of embedding strategies. For each MP term, sets of 15 nearest neighbors were computed in all embeddings. The similarity of neighborhoods were computed using the Jaccard index. The similarity for a pair of embeddings was defined as the mean Jaccard index for all MP terms. All approaches were analyzed in duplicate (two embeddings produced with different seeds for random number generators) and compared with a random layout with MP terms arranged uniformly at random in two dimensions.

**Figure S2.**
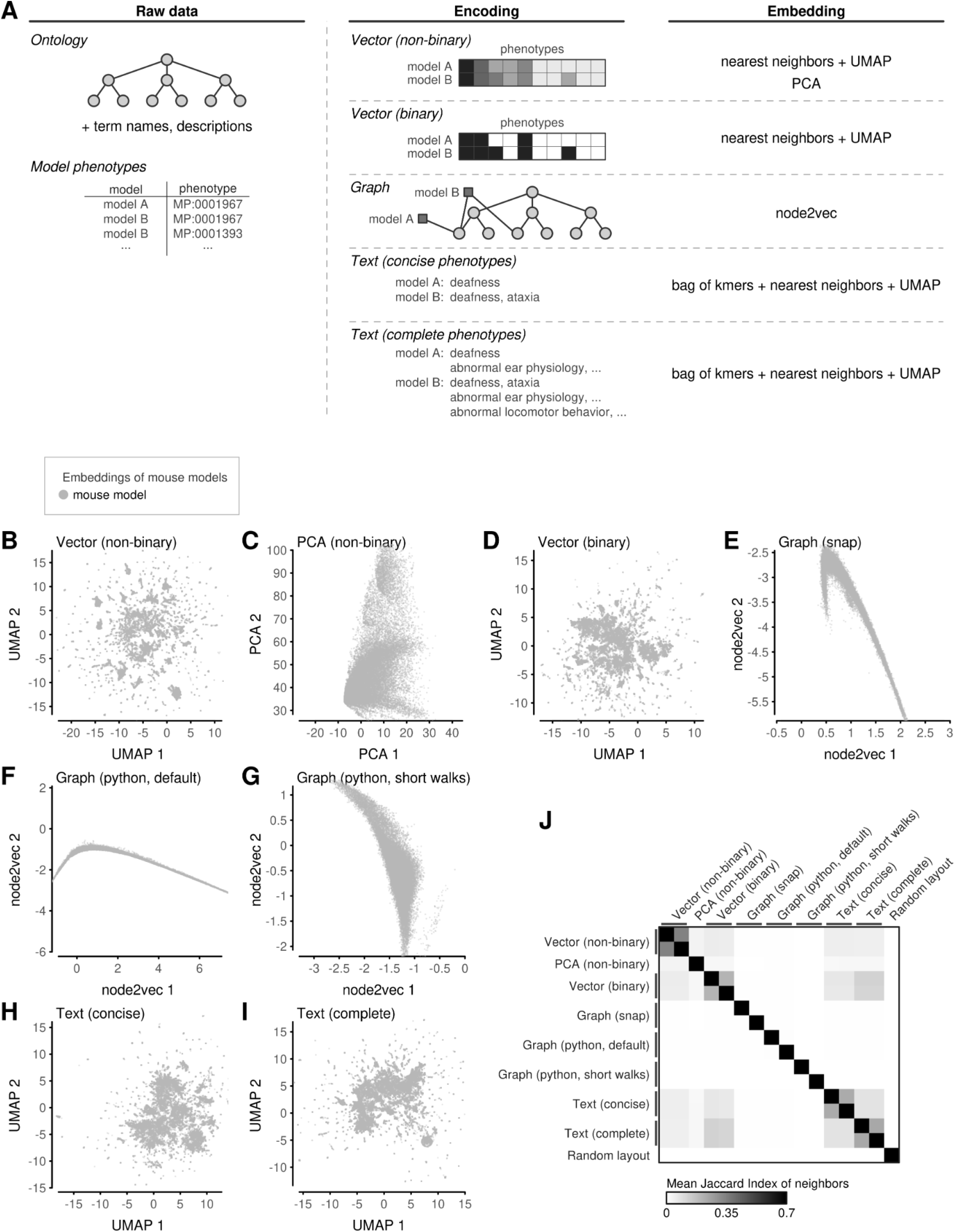
Embeddings of mouse models. (A) Schematic of phenotype data for mouse models, possibilities for data encodings, and algorithms for creating embeddings. (B) UMAP embedding based on non-binary vector representations. (C) PCA based on non-binary vector representations. (D) UMAP embedding based on binary vector representations. (E) Node2vec embedding based on a graph connecting mouse models to their ontology phenotypes, generated using snap implementation of node2vec with default settings. (F) Similar to (E), generated using python implementation of node2vec with default settings. (G) Similar to (E), generated with python implementation of node2vec with settings walk-length and num-walks set to 5. (H) Embedding based on text descriptions of mouse phenotypes. (I) Similar to (H), but using text descriptions of complete phenotypes. (J) Comparison of embedding strategies. For each mouse model, sets of 15 nearest neighbors were computed in all embeddings. The similarity of neighborhoods were computed using the Jaccard Index. The similarity for a pair of embeddings was defined as the mean Jaccard index for all mouse models. All approaches were analyzed in duplicate (two embeddings produced with different seeds for random number generators) and compared with a random layout with mouse models arranged uniformly at random in two dimensions.

**Figure S3.**
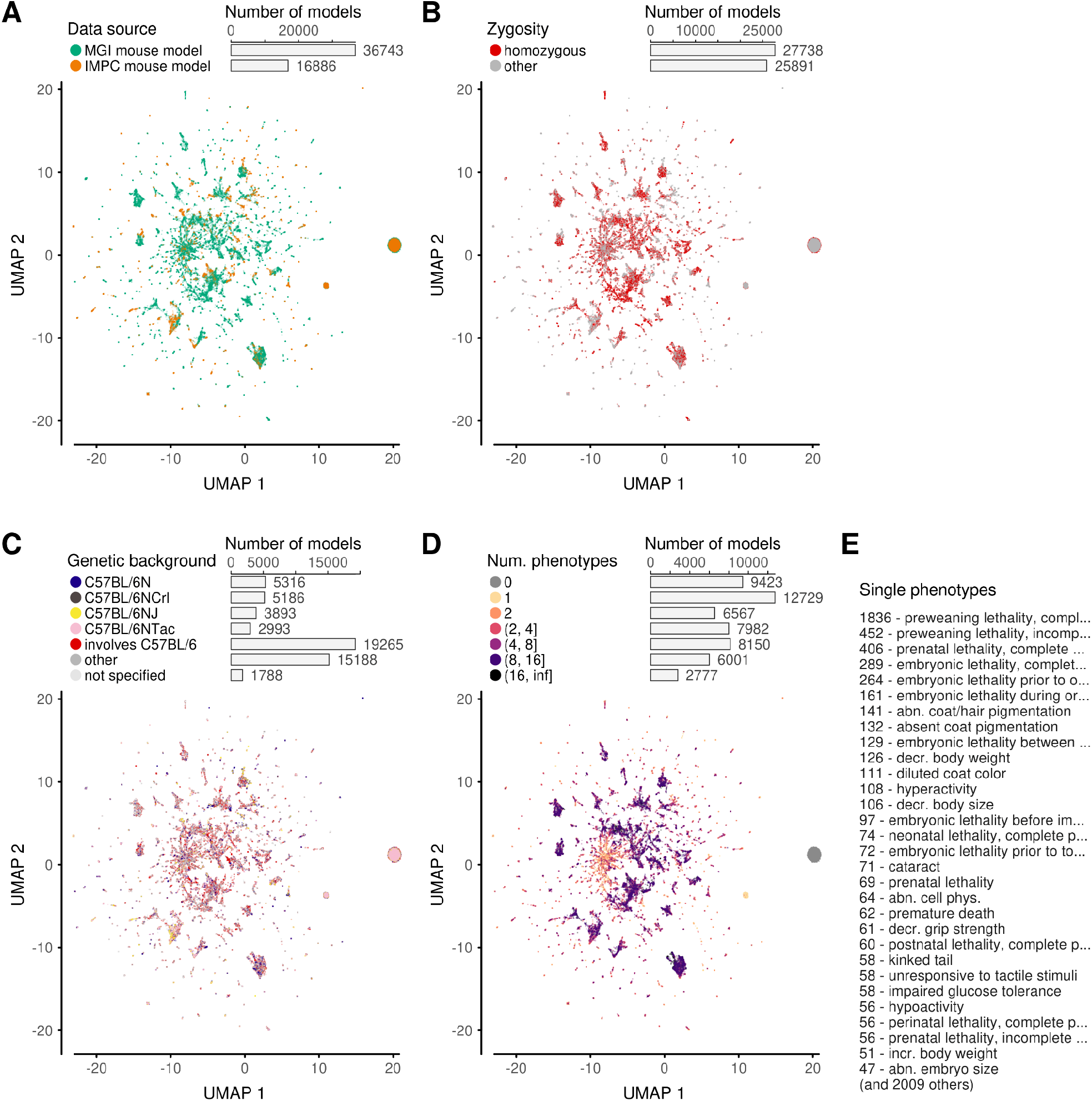
Mouse model covariates. Embeddings of mouse models based on non-binary vectors of phenotypes. Four panels differ by stratification strategy: (A) phenotyping source; (B) zygosity of gene knock-out; (C) animal genetic background; (D) number of phenotypes. (E) Listing of the most frequent MP terms present among models with a single phenotype. Leading numbers indicate the number of models that are annotated only with the stated term.

**Figure S4.**
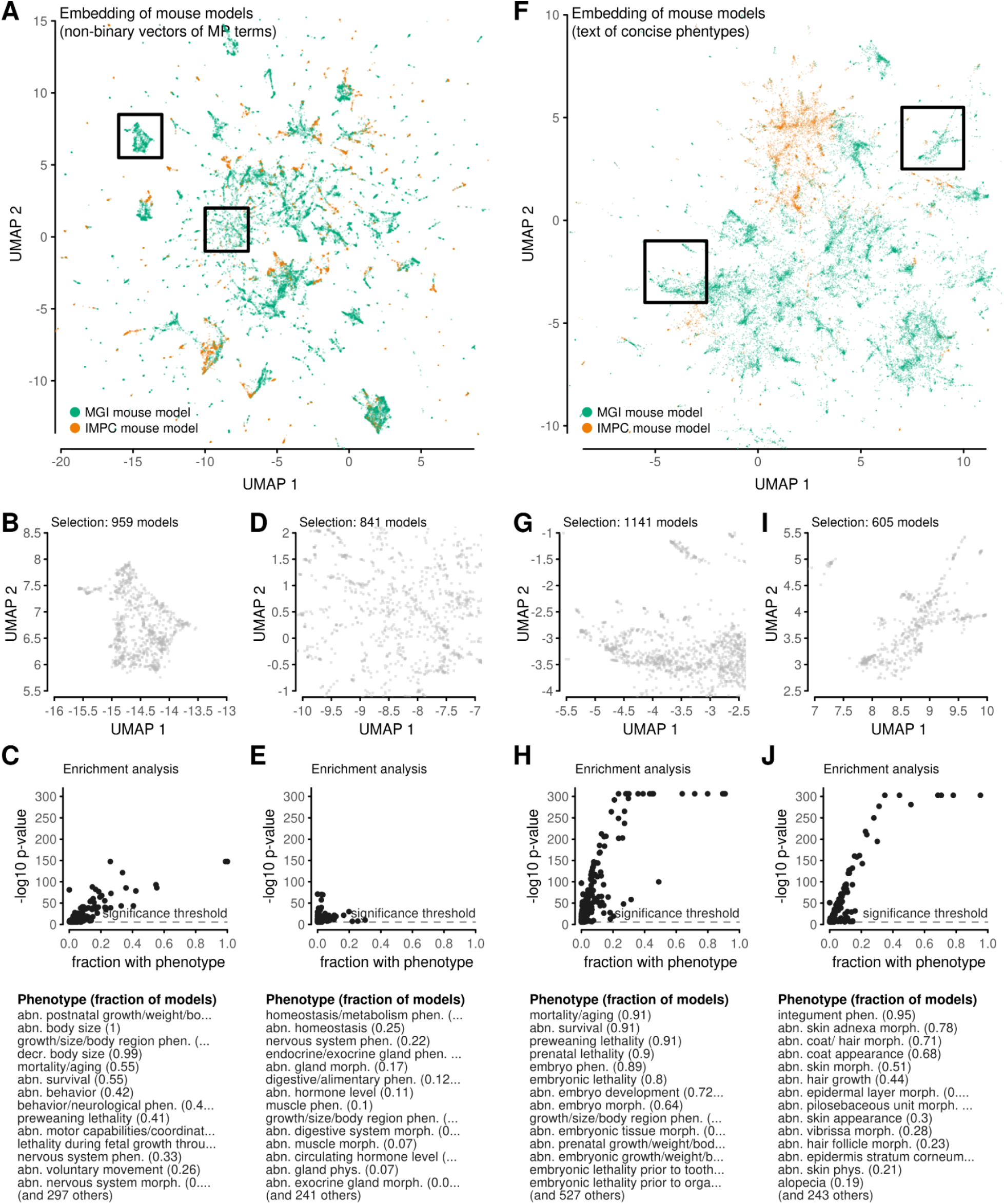
Feature enrichment in embedding regions. (A) Embedding of all mouse models based on vector representations of phenotypes. Two regions are selected with rectangles. (B) A detailed view of one of the selected regions from (A). (C) Enrichment analysis comparing phenotypes observed in animal models shown in (B) compared to all other models outside the selected region. Dots correspond to MP phenotypes. Axes show the fraction of selected models with a given phenotype, and an enrichment significance score for the phenotype computed using a Fisher test (some p-values are truncated). Significance level is p=0.05 after Bonferroni correction. The table at the bottom names the most significant phenotypes. (D, E) Analogous to (B, C), but treating another region. (F-J) Analogous to (A-E), but based on an embedding of mouse models based on concise text descriptions.

**Figure S5.**
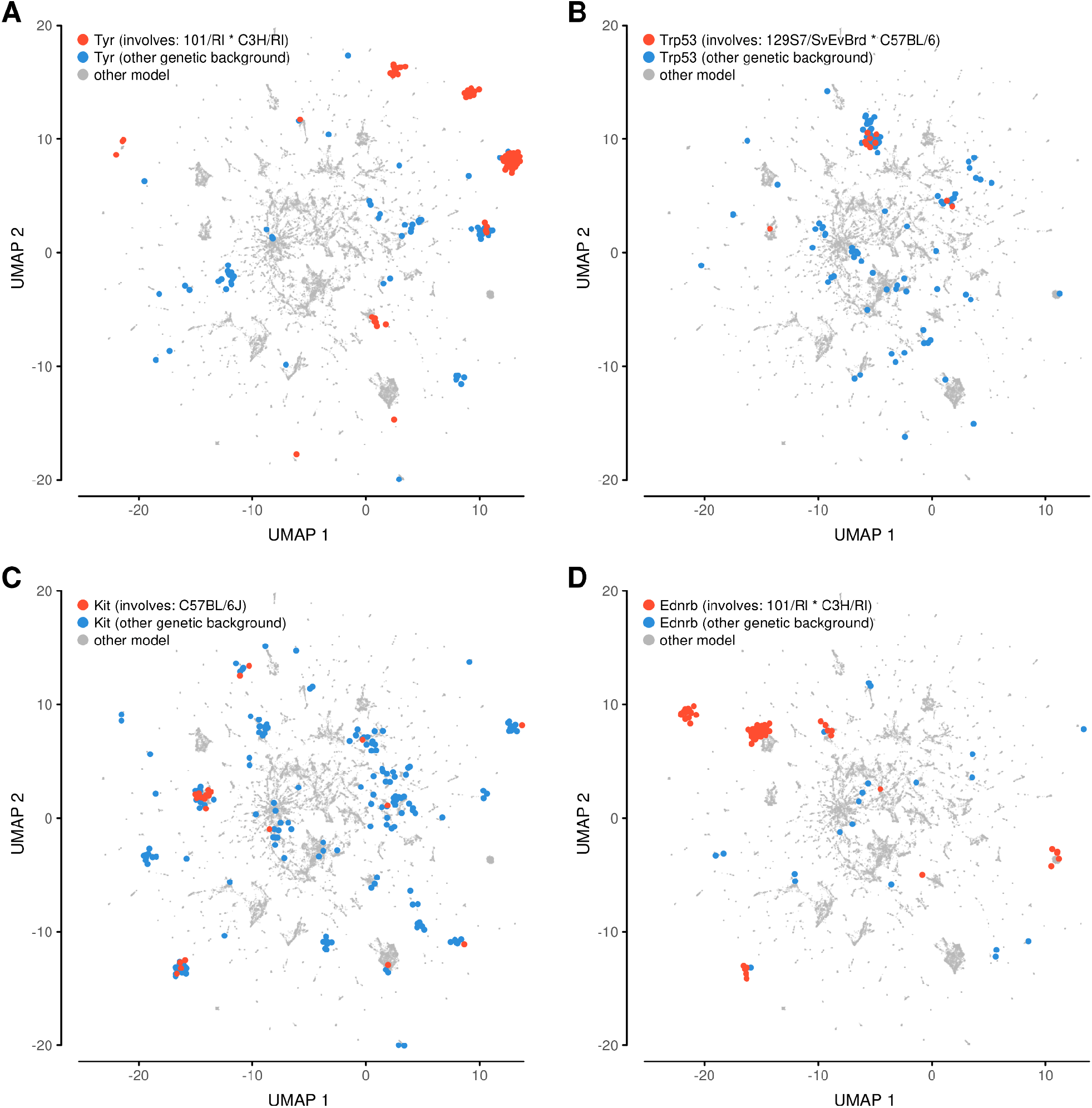
Effect of genetic background for selected genes. All panels show embeddings of mouse models based on non-binary vectors. In each panel, models with a single gene are highlighted: (A) Tyr; (B) Trp53; (C) Kit; and (D) Ednrb. One set of highlighted models reveals the most common genetic background (for that gene). The other highlighted set includes models with other genetic backgrounds (for that gene). The position of highlighted models are jittered to better reveal the number of models in dense areas.

**Figure S6.**
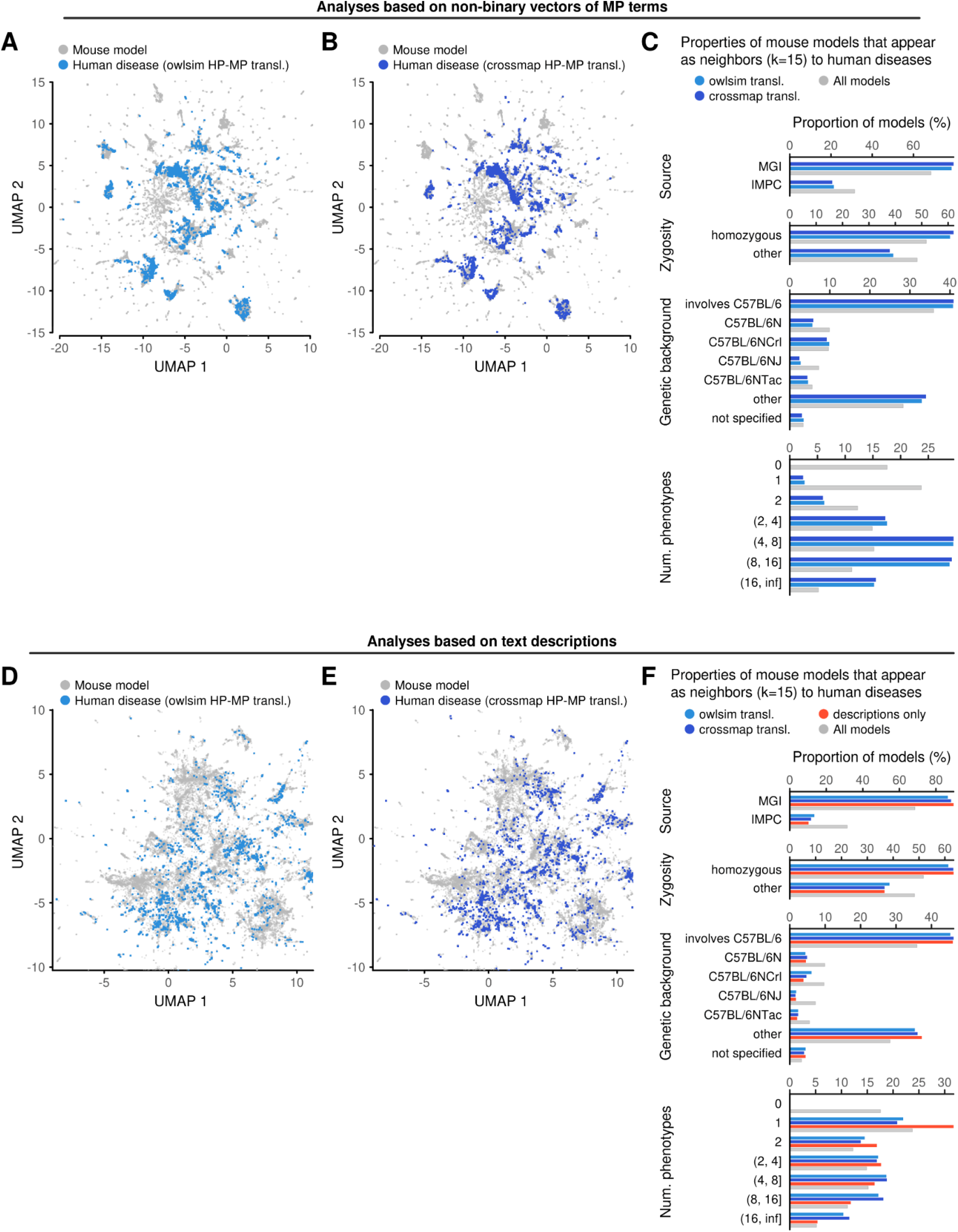
Analyses of diseases. (A) Projection of human disease phenotype profiles (colored points) into an embedding of mouse models (gray dots). The embedding was created based on non-binary vectors with mouse model phenotypes, without information about diseases. Disease phenotype profiles were translated from the human phenotype (HP) into the mammalian phenotype (MP) ontology terms using owlsim, encoded into vectors, and projected into the embedding using UMAP. (B) Similar to (A), but with the translations between human and mammalian ontologies carried out using crossmap. (C) Comparison of the properties of mouse models identified during disease analysis to the properties of the entire mouse model cohort. Disease profiles were compared with all mouse models, recording 15 nearest mouse models for each disease. The set of thus selected models were stratified according to the mouse model data source, mutation zygosity, mouse strain/genetic background, and the number of phenotypes annotated to the mouse models. The same stratification was applied to the entire set of mouse models for comparison. (D, E, F) Analogous to panels (A, B, C), but with all analyses carried out using text-based similarities. Analysis in (F) includes a group of models identified during analysis of diseases that have text descriptions but are not annotated with formal ontology-based phenotypes.

